# Two strains of Toscana virus show different virulence and replication capacity in mice and cell culture models

**DOI:** 10.1101/2024.12.11.627885

**Authors:** Marlène Roy, Sandra Lacôte, Sophie Desloire, Adrien Thiesson, Coralie Pulido, Noémie Aurine, Cyrille Mathieu, Bertrand Pain, Philippe Marianneau, Frédérick Arnaud, Maxime Ratinier

## Abstract

Toscana virus (TOSV), belonging to *Phenuiviridae* family, is circulating in most Mediterranean countries and is transmitted to humans by infected female sand flies. While most infections are asymptomatic, TOSV is considered as a leading cause of meningitis and encephalitis in humans during summer. Three TOSV genotypes (named A, B, and C) have been identified, although no virus strain belonging to lineage C has been isolated so far. To date, the relationship between TOSV genetic diversity and viral pathogenicity or replication capacity remains unknown. This study aimed to compare two TOSV strains from either lineage A (TOSV-A) or B (TOSV-B) in several cell culture and two mouse models. We showed that TOSV-A replicated more efficiently in BSR and A549 cells while TOSV-B had a replication advantage in human induced pluripotent stem cells differentiated in neural cells and LL-5 sand fly cells. *In vivo*, we were unable to detect any virus in the brains of immunocompetent C57BL/6JRj mice infected with either strain of TOSV. On the contrary, we showed that TOSV-A disseminated to the central nervous system of 129/Sv *ifnar* ^-/-^ mice unlike TOSV-B, despite higher viremia of TOSV-B and a greater dissemination of this strain in other organs. The reasons for these differences are not yet known, although we showed that the presence of TOSV neutralizing antibodies in serum was slightly delayed in TOSV-A-infected mice. Altogether, the data presented in this study provide new avenues to study TOSV-induced pathogenesis and ultimately unveil molecular viral determinants modulating TOSV replication capacity.

**Author Summary:** Toscana virus is a sand fly-borne phlebovirus with tropism for the human central nervous system (CNS) and the causal agent of aseptic meningitis and encephalitis during summer in countries surrounding the Mediterranean Sea. The pathogenesis induced by TOSV is still poorly understood as well as the correlation between TOSV genetic diversity and viral pathogenicity or replication capacity. In this study, we characterised two TOSV strains belonging to two different genetic lineages (named A and B) in several *in vitro*, *ex vivo* and *in vivo* models of infection. We showed that the biological properties of these two strains is strikingly different and, importantly, we identified 129/Sv *ifnar* ^-/-^ mice as a suitable model to study the mechanism of TOSV invasion of the CNS and viral pathogenesis. Our comparative study also highlights the importance of strengthening our efforts to identify molecular viral determinants modulating TOSV-induced pathogenesis and vector transmission, in order to better characterise the epidemiological risk posed by circulating strains.

## Introduction

Toscana virus (TOSV) has been isolated for the first time in 1971 from Phlebotomine sand flies in Italy [1]. It is circulating in many countries surrounding the Mediterranean Sea and is transmitted to human by infected female sand flies [2]. Since its discovery in *Phlebotomus perniciosus and P. perfiliewi*, many other sand fly species have been tested positive for TOSV, such as *P. longicuspis*, *P. sergenti*, *P. tobbi* and *Sergentomyia minuta* [3–5]. In humans, TOSV infection is mostly asymptomatic [6]. The predominant symptomatic form is characterized by fever, headache, and nausea, with low and transient viremia [7,8]. However, TOSV can also invade and damage the central nervous system (CNS) and is responsible for aseptic acute meningitis, meningoencephalitis, and encephalitis, with an incubation period estimated at 12 days post-infection [9,10]. Seroprevalence in human population varies widely by country/region and is highly dependent on the age of tested population as well as the abundance and exposure to the vector [11–15].

TOSV is a member of the Sandfly fever Naples virus (SFNV) complex and belongs to the genus *Phlebovirus* within the *Phenuiviridae* family, order *Hareavirales* [16]. TOSV has a tri-segmented single-stranded RNA genome of negative or ambisense polarity [17,18]. The Large segment (called L) encodes the viral RNA dependent RNA polymerase (RdRp). The Medium segment (called M) encodes a polyprotein cleaved into the non-structural protein of M segment (NSm) and the two glycoproteins Gn and Gc. The Small segment (called S) encodes the nucleoprotein (N, positive sense) and the non-structural protein of S segment (NSs, negative sense), an antagonist of type I interferon response [19,20]. Phylogenetic analysis of TOSV genome revealed three genetically distinct lineages (called A, B, and C) [9]. To note, lineage C has been identified in Croatia and Greece, but no virus has been isolated until now [21,22]. Lineage A has been shown to circulate in Italy, Romania, Tunisia, and Algeria whereas lineage B has been found in Spain, Portugal, and Morocco [23–28]. Several lineages may circulate in the same geographical area, such as lineages A and B co-circulating in France and Turkey, or lineages B and C co-circulating in Croatia [29–31]. To date, no clear evidence of reassortment between different lineages has been demonstrated for TOSV.

The pathogenesis of TOSV is still poorly understood, not only because TOSV neurological infection is underdiagnosed but also due to the difficulty to develop an animal model. Indeed, it has been shown that TOSV strain 1812 (lineage A) was not virulent in newborn or adult mice even when injected intracerebrally (i.c.) [32]. Only the strain 1812V obtained by serial passaged in Vero and McCoy cells as well as in brain of newborn mice induced the death of the adult BALB/c mice between day 6 and day 10 post-infection, when inoculated i.c.. In these conditions, immunohistochemical analysis shows that this neuro-adapted strain replicates efficiently in neurons and induced apoptosis.

To date, it is unknown how viral genetic diversity is affecting the virus-host-vectors interactions. In this study, we characterized two TOSV strains from either lineage A (TOSV-A) or lineage B (TOSV-B) in several *in vitro, ex vivo*, and *in vivo* models. We showed that the replication capacity of TOSV is highly dependent on the strains considered as well as the *in vitro* cell culture models used. Moreover, we showed that immunocompetent C57BL/6JRj mice rapidly resolved the infection by both TOSV strains and that no virus was detectable in the brain. In contrast, 129/Sv *ifnar* ^-/-^ mice were susceptible to TOSV infection, with TOSV-A showing a strong tropism for the CNS, unlike TOSV-B infection that led to a higher viremia with an efficient dissemination to other organs. The difference of virulence observed between the two TOSV strains is still not understood but we showed that the appearance of neutralizing antibodies (nAbs) is slightly delayed in TOSV-A-infected animals compared to TOSV-B, and that TOSV-B is able to replicate actively in human-induced pluripotent stem cells-derived neural cells as well as in mice organotypic brain cultures. Altogether, the data presented in this study provide a solid basis for further study of TOSV-induced pathogenesis and demonstrate that 129/Sv *ifnar* ^-/-^ mice are a relevant model of TOSV infection.

## Materials and Methods

### Ethics statement

The experimental protocols complied with the regulation 2010/63/CE of the European Parliament and of the Council of 22 September 2010 on the protection of animals used for scientific purposes and as transposed into French law. These experiments were approved by the Anses/ENVA/UPEC ethics committee and the French Ministry of Research (Apafis n° 2017110715154626 (#17084)). All *ex vivo* experiments with mice were performed by CM (accredited by the French veterinary service) according to the French National Charter on the ethics of animals. Briefly, mice were directly euthanized by decapitation according to the American Association for Accreditation of Laboratory Animal Care (AAALAC) recommendations and according to French Ethical Committee (CECCAPP) regulations (accreditation # CECCAPP_ENS_2014_034), before preparation of organotypic brain cultures.

### Cell culture

VeroE6, BSR (BHK21 clone, kindly provided by Karl K. Conzelmann) [33], and A549 (kindly provided by Richard E. Randall, University of St Andrews, UK) cells were grown in Dulbecco’s Modified Eagle Medium (DMEM, Gibco, Thermo Fischer Scientific, Villebon-sur-Yvette, France) supplemented with 10% heat-inactivated fetal bovine serum (HI-FBS, GE HEALTHCARE Europe GmbH, Freiburg, Germany). Human-induced pluripotent stem cells (hiPSCs), declared onto the CODECOH platform with the number DC-2021-4404, were obtained from two different donors and subsequently differentiated into neurons, astrocytes, and oligodendrocytes for 45 days as previously described [34]. Mammalian cell lines were cultured at 37°C in a 5% CO_2_ humidified atmosphere. LL-5 cells (established from sandfly *Lutzomyia longipalpis* embryo; kindly provided by Robert Tesh, University of Texas Medical Branch, US) were cultivated in Schneider’s insect medium (Gibco, Thermo Fischer Scientific) supplemented with 20% HI-FBS. Insect cells were incubated at 28°C without CO_2_.

### Viruses

TOSV strain 1500590 (Lineage A, referred as TOSV-A) was isolated in Vero cells from human patient serum [35]. TOSV strain MRS2010-4319501 (Lineage B, referred as TOSV-B, obtained from the European Virus Archive collection) was isolated in Vero cells from the cerebrospinal fluid of a human case with meningitis [36]. Both strains were isolated in Marseille, France. TOSV stocks were generated by infecting BSR cells and virus titres were determined by plaque-forming assay using BSR cells, as already described [37].

### Virus growth curves

hiPSCs-derived neural cells were infected with 2x10^4^ plaque-forming units (PFU) of either TOSV-A or TOSV-B, corresponding to an approximated multiplicity of infection (MOI) of 0.1. LL-5, BSR, and A549 cells were infected with either TOSV-A or TOSV-B at a MOI 0.01. Cell supernatants were collected at 24-, 48-, and 72-hours post-infection (hpi). The amount of infectious viral particles was determined by endpoint dilution assay using BSR cells. Viral titres were calculated by Reed and Muench’s method and expressed as tissue culture infectious dose 50 per millilitre (TCID_50_/ml) [38]. Experiments with hiPSCs-derived neural cells were performed twice in triplicate, each time with cells from two different donors (n=12). Experiments with LL-5, BSR, and A549 cells were performed three times independently and in triplicate (n=9). All virus growth curves were obtained using two different stocks of each virus. Statistical analyses were conducted at each time point using Mann-Whitney test within Graphpad Prism 8.4 (Graphpad Software Inc., La Jolla, CA, USA).

### *In vivo* studies

#### Study 1

Six-8 weeks old C57BL/6JRj wild type (n=12, two independent experiments) and 129/Sv *ifnar* ^-/-^ (n=18, three independent experiments) female mice were subcutaneously (SC) infected with 10^3^ PFU of either TOSV-A or TOSV-B. Mock groups contained six or ten animals, respectively. Mice were euthanized at the end of the course of the experiment (13 days post-infection, dpi) or earlier if the animal showed severe clinical signs. Brains and livers were therefore collected and tested for the presence of viral RNA by RT-qPCR and, if positive, the amount of infectious viral particles was determined by plaque-forming assay using VeroE6. Survival curves were analysed using Graphpad Prism 8.4 and statistical analyses were conducted using Log-rank (Mantel-Cox) test. Sera were also collected on the day of the infection (day 0) and then at 3, 5, 7, 10, and 13 dpi. We analysed: (i) the production of anti-TOSV IgM and IgG using enzyme-linked immunosorbent assay (ELISA), (ii) the presence of viral RNA by RT-qPCR and (iii) in case of TOSV RNA detection (129/Sv *ifnar* ^-/-^ mice only), the infectious viral titre by plaque-forming assay using VeroE6.

#### Study 2

6-8 weeks old C57BL/6JRj wild type (n=4) and 129/Sv *ifnar* ^-/-^ (n=4) female mice were SC-infected with 10^3^ PFU of either TOSV-A or TOSV-B. Mice were euthanized at either 3 (n=2) or 5 dpi (n=2). Organs (liver, heart, lung, brain, spleen, lymph nodes and kidney) and sera were collected, the presence of TOSV RNA was assessed using RT-qPCR and, if positive (129/Sv *ifnar* ^-/-^ mice only), the infectious viral titre (expressed as PFU/g for tissues or PFU/ml for sera) was determined using plaque-forming assay using VeroE6.

### RT-qPCR

Brain, liver, heart, lung, brain, spleen, lymph nodes, and kidney of euthanized mice were weighed and homogenized 3 times for 30 s at 30 Hz using Tissue Lyser II (Qiagen, Courtaboeuf, France) in 500 µl of DMEM with 2 stainless steel beads (Thermo Fisher Scientific, Villebon-sur-Yvette, France). Samples were subsequently centrifugated at 500 g for 5 min and supernatants were collected. Viral RNA was isolated from supernatants or sera using QIAamp Viral RNA Mini Kit (Qiagen) and the QIAcube (Qiagen) automate, according to the manufacturer’s protocol. One step reverse transcription quantitative qPCR was performed using SuperScript™ III One-Step RT-PCR System with Platinum™ Taq DNA Polymerase (Invitrogen by Life Technologies, Thermo Fisher Scientific) according to the manufacturer’s instructions. Quantitative PCR was performed with the primers STOS-50F (5’-TGC TTT TCT TGA GTC TGC AG -3’), STOS-138R (5’-CAA TGC GCT TYG GRT CAA A -3’), and the TaqMan® probe STOS-84T-FAM (FAM 5’ – ATC AAT GCA TGG GTR AAT GAG TTT GC TAC C-3’ TAMRA) [39]. The RT-qPCR cycling conditions were 45°C for 30 min, 95°C for 10 min followed by 45 cycles set up as follow: 95°C for 2 s and 60°C for 20 s.

### ELISA

Anti-TOSV antibodies within mice sera were detected using in-house IgM and IgG ELISAs. TOSV antigens (Ag) were prepared from VeroE6 cells infected with TOSV-A strain (MOI=0.01; 3 days post-infection) and negative Ag were obtained from non-infected VeroE6 cells, as previously described (34).

For IgM detection, 96 -well plates (Nunc Maxisorp, Thermo Fisher Scientific) were coated the day before with rabbit anti-mouse IgM antibody (100 μL/well, 1:500 dilution, SAB3701197, Sigma-Aldrich, Merck, Saint-Quentin Fallavier, France). Then, they were incubated with 100 μL/well of 1:100 dilution of mice sera. TOSV or negative Ag (1:8 dilution) were then added to the plate and were subsequently detected with hyper-immunised sera from hamsters infected with TOSV-A strain (1:1000 dilution) and finally by goat anti-Hamster IgG (H+L)-Horseradish Peroxidase (HRP) conjugated antibody (1:3000 dilution, Life Technology, Thermo Fisher Scientific).

For IgG detection, plates were coated with TOSV or negative Ag (1:500 dilution), further incubated with 100 μL/well of 1:100 dilution of mice sera and subsequently with HRP-conjugated rabbit anti-mouse IgG (whole molecule, 1:5000, Sigma-Aldrich).

HRP enzymatic activity was determined using TMB substrate (Thermo Fisher Scientific) and stopped with 10.6% phosphoric acid solution. Optical density at 450 nm (OD450) was measured using a TECAN microplate reader. Corrected OD450 (ODc450) was calculated by subtracting OD450 from negative Ag to OD450 from TOSV Ag. Sera were considered positives when ODc450 values were >0.1.

### Virus neutralization test (VNT)

The sera positive for anti-TOSV IgG and/or IgM (from one independent experiment only, n=6) were tested for the presence of nAbs against TOSV by in-house VNT. Dilutions of each serum were mixed with a viral dilution containing 100-200 plaque-forming units of TOSV-A and were incubated 1 hour at 37°C in a humid atmosphere. The mix were then added to confluent monolayer of VeroE6 cells in 96 wells plates. After an incubation of 1 hour at 37°C, the cultures were overlaid with fresh DMEM medium. After 5 days of incubation at 37°C, the cells were stained with crystal violet solution. The neutralizing titre was determined as the highest dilution of serum that caused complete cytopathic effect.

### Organotypic brain cultures

Organotypic brain cultures (OBC) were prepared from C57BL/6 and *ifnar* ^-/-^ mice (7 days old) and cultured as already described [40]. Each OBC was infected with 10^3^ PFU of either TOSV-A or TOSV-B and subsequently collected at 24-and 72-hpi. OBC were subsequently crushed in 300 µl PBS with a tungsten carbide bead using the TissueLyser II (Qiagen) and centrifugated to remove the cellular debris. 100 µl of the clarified supernatant was used to extract total RNA and perform RT-qPCR as already described. The remaining supernatant was used to assess viral titres by endpoint dilution assay as described above. Each experiment was performed twice independently using two different virus stocks. Statistical analysis was conducted at each time point using Mann-Whitney test within Graphpad Prism 8.4.

## Results

### Replication kinetics of TOSV-A and TOSV-B

Since TOSV shows a strong tropism for CNS in severe forms of the disease, we first analyzed the replication kinetics of TOSV-A and TOSV-B in hIPSc-derived neural cells (Fig 1).

**Figure 1:**
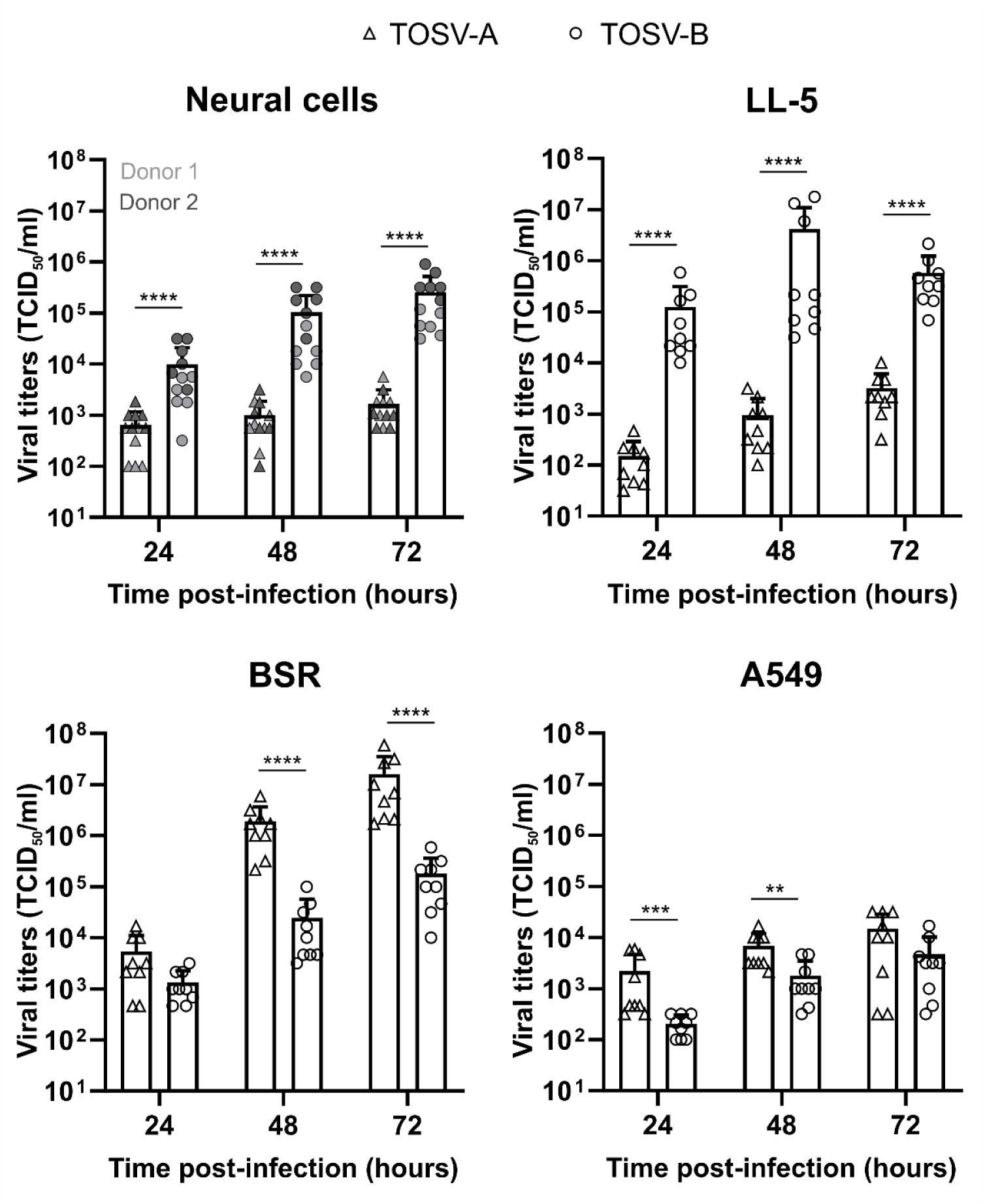
Growth properties of TOSV-A and TOSV-B. Viral growth curves in hIPSc-derived neural cells (MOI ≈ 0.1; 2x10^4^ PFU), LL-5, BSR, and A549 cells (MOI: 0.01) infected by either TOSV-A or TOSV-B. Supernatants were harvested at 24, 48, and 72 hpi, titrated by endpoint dilution and viral titers expressed as TCID_50_/ml. Comparisons of viral titers at each time point between TOSV-A and TOSV-B were carried out using the Mann-Whitney test; P > 0.05 (ns), P < 0.01 (**), P < 0.001 (***), and P < 0.0001 (****).

TOSV-A replication was significantly slower than TOSV-B with titers approximately 15-(24 hpi) and 150-fold (72 hpi) lower. Moreover, TOSV-B titers were increasing through the time to reach 2.5x10^5^ TCID_50_/ml at 72 hpi whereas TOSV-A infectious viral particles production is stable at around 10^3^ TCID_50_/ml. This phenotype is observed for both donors, although TOSV-B titers were higher in hIPSc-derived neural cells from donor 2 for which cell density was more important. Next, we compared the growth properties of the two TOSV strains in LL-5 cells obtained from embryo of sand fly *Lutzomyia longipalpis*. TOSV-B consistently reached higher titers (2.2 to 3.6 log_10_-fold increase) than TOSV-A, with peak titers at 48 hpi (4.2x10^6^ TCID_50_/ml) and 72 hpi (3.2x10^3^ TCID_50_/ml), respectively. On the contrary, TOSV-A grew to higher titers in BSR (which are interferon incompetent) and A549 (which are interferon competent) cells compared to TOSV-B. Both strains reached higher titers in BSR (1.6x10^7^ and 1.8x10^5^ TCID_50_/ml at 72 hpi for TOSV-A and TOSV-B, respectively) than in A549 (1.5x10^4^ and 4.7x10^3^ TCID_50_/ml at 72 hpi), likely because there is no interferon response to overcome in BSR cells. Overall, these data showed that the growth properties of the two TOSV strains are different and highly depend on the infected cell types.

### C57BL/6JRj mice are poorly infected by both TOSV-A and TOSV-B and do not develop disease symptoms

We next SC-infected C57BL/6JRj female mice with 10^3^ PFU of either TOSV-A or TOSV-B to investigate their virulence in immune-competent mice. Only one mouse in TOSV-A group died at 7 dpi when no mice succumbed in TOSV-B group (Fig 2A). The dead mouse and all the mice euthanized at the end of the experiment in the two groups were negative by RT-qPCR for TOSV RNA in their liver and brain. Moreover, we detected very low level of TOSV RNA in the serum (RNAemia) of one mouse at 3 dpi, three at 5 dpi and two at 7 dpi in TOSV-B group, whereas no mice showed RNAemia in TOSV-A group (Fig 2B and S1A). To note, no infectious viral particle was detected by plaque-forming assay in the sera of the mice with RNAemia. In parallel, we also investigated the dynamics of IgM and IgG seroconversion in TOSV-infected mice (Fig 2C and 2D, respectively). All the mice infected by TOSV-B produced specific TOSV IgM at 5 dpi and IgG at 10 dpi. By comparison, in the group of TOSV-A-infected mice, four mice out of seven (57%, 10 dpi) and six mice out of ten (60%, 13 dpi) showed specific TOSV IgM response, when only two mice out of eight (25%, 10 dpi) and two mice out of ten (20%, 13 dpi) produced TOSV-specific IgG. Additionally, nAbs titers measured by VNT in sera of TOSV-B-infected mice (n=6) were increasing from 6.6 log_2_ (VNT titer) at 5 dpi to 11.6 log_2_ (VNT titer) at 13 dpi, when only one mouse infected with TOSV-A produced nAbs at 5 and 7 dpi (the one that died at 7 dpi) and another one at 10 and 13 dpi (Fig 2E). The latter had VNT titers lower than the one observed in TOSV-B-infected mice.

**Figure 2:**
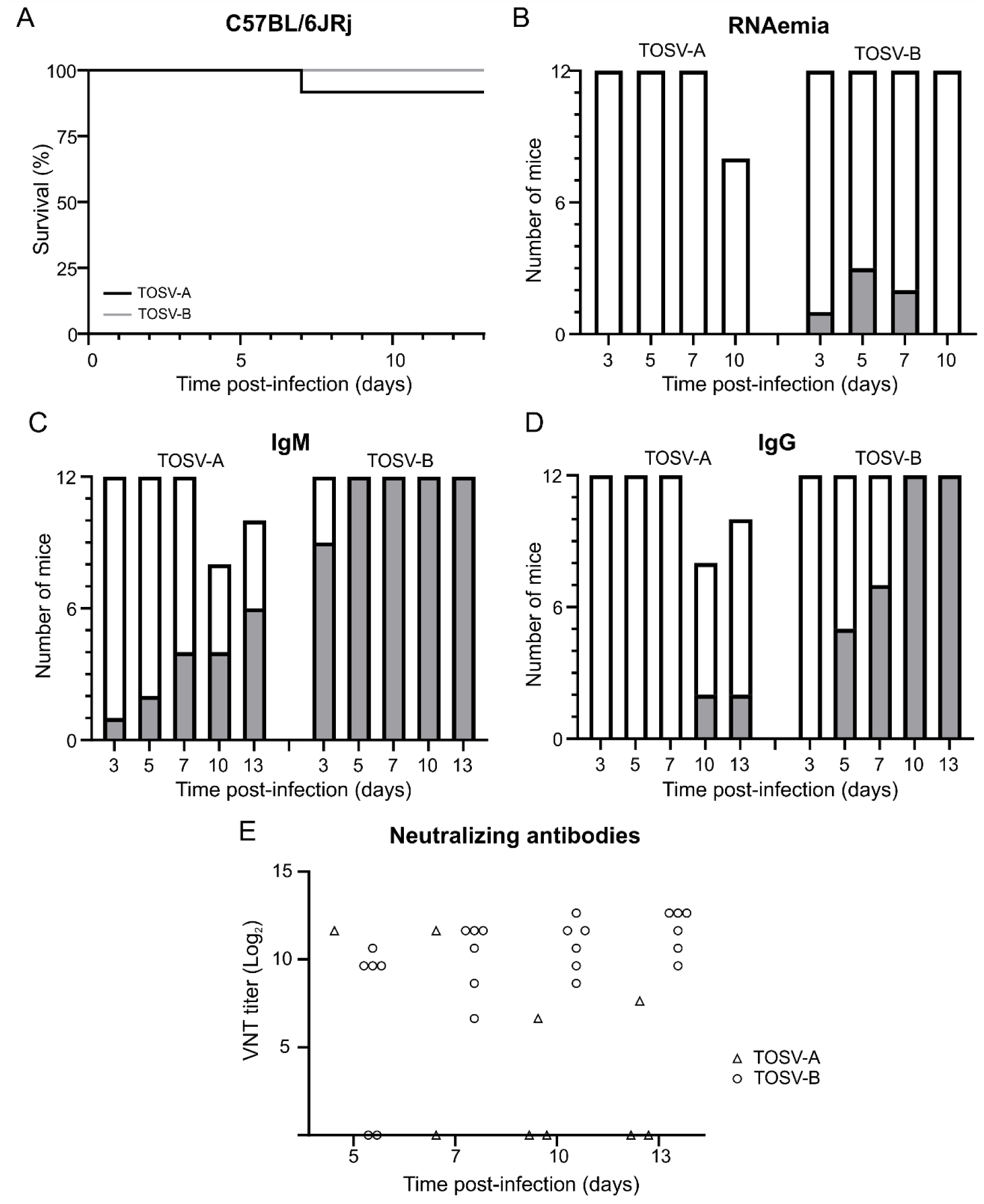
Experimental infection of C57BL/6JRj mice with TOSV-A and TOSV-B. (A) Kaplan-Meier survival curves. 6-8 weeks old female C57BL/6JRj mice (n=12) were infected subcutaneously with 10^3^ PFU of either TOSV-A or TOSV-B. Survival curves were analysed using Log-rank (Mantel-Cox) test. (B) Detection of TOSV RNA in the sera (RNAemia). Sera were collected at 3, 5, 7, and 10 dpi. Levels of viral RNA in sera of infected mice were measured by RT-qPCR targeting segment S. Positive mice sera are represented in gray and negative mice sera are represented in white. Note that only 8 mice were analysed at 10 dpi for TOSV-A group because the sera of three mice have not been collected and one mouse died suddenly at 7 dpi. Kinetics of IgM (C) and IgG (D) antibody response against TOSV-A and TOSV-B. Sera were collected at 3, 5, 7, 10, and 13 dpi. Anti-TOSV antibodies within mice sera were detected using in-house IgM and IgG ELISAs. At the indicated day, grey bars represent seroconverted mice and white bars mice with non-detectable anti-TOSV IgM or IgG antibodies. Note that only 10 mice were analysed at 13 dpi for TOSV-A group because the sera of one mouse was not collected and one mouse died suddenly at 7 dpi. (E) Virus neutralisation test. Sera (from one independent experiment only, n=6) containing anti-TOSV IgG and/or IgM were tested for the presence of nAbs against TOSV. The neutralizing titre, expressed as log_2_, was determined as the highest dilution of serum that caused complete cytopathic effect.

Next, we aim to refine the viral replication kinetics and organ tropism at early time points after inoculation of the two TOSV strains. To do so, we infected C57BL/6JRj female mice with either TOSV-A or TOSV-B and collected sera and organs (liver, heart, lungs, brain, spleen, lymph nodes, and kidney) at 3 and 5 dpi (n=2 for each group). Sera and organs of TOSV-A-infected mice were all negatives for TOSV RNA, when one mouse infected with TOSV-B had TOSV RNA in the liver, heart, lungs and lymph nodes at 3 dpi (Fig S1B). Interestingly, the spleen of all mice infected by TOSV-B were positive for TOSV RNA at both 3 and 5 dpi. Overall, Ct values were quite high and range between 30 and 36 (Fig S1B), highlighting the low amount of viral RNA in the samples. Altogether, these data showed that C57BL/6JRj mice are resistant to both TOSV strains infection when inoculated subcutaneously. Therefore, this mouse model of infection is not relevant for studying TOSV-induced pathogenesis.

### 129/Sv *ifnar*^-/-^ mice are susceptible to TOSV infection and TOSV-A strain is more virulent than TOSV-B strain

The interferon (IFN) pathway is often considered as the first line of defense against viral infection and, therefore, is crucial to control viral replication and dissemination. Since TOSV replication observed in C57BL/6JRj mice was low, especially for TOSV-A strain, we SC-infected 129/Sv *ifnar* ^-/-^ mice, unable to respond to type I IFNs, with 10^3^ PFU of either TOSV-A or TOSV-B. The Kaplan-Meier curves showed that TOSV-A strain was significantly more virulent than TOSV-B (P= 0.0419, Log-rank (Mantel-Cox) test; Fig 3A). Indeed, eight mice succumbed from TOSV-A infection between 8 and 11 dpi (survival rate of 55.6%), when two mice died in TOSV-B group at 8 dpi (survival rate of 88.9%) (Fig 3A). Interestingly, as for C57BL/6JRj mice, more 129/Sv *ifnar* ^-/-^ mice infected with TOSV-B have detectable TOSV RNA in the sera (RNAemia) than those infected by TOSV-A (Fig 3B and S1B). For both groups, RNAemia was mainly observed between 3 and 7 dpi, with fewer mice tested positive at 8-10 dpi.

**Figure 3:**
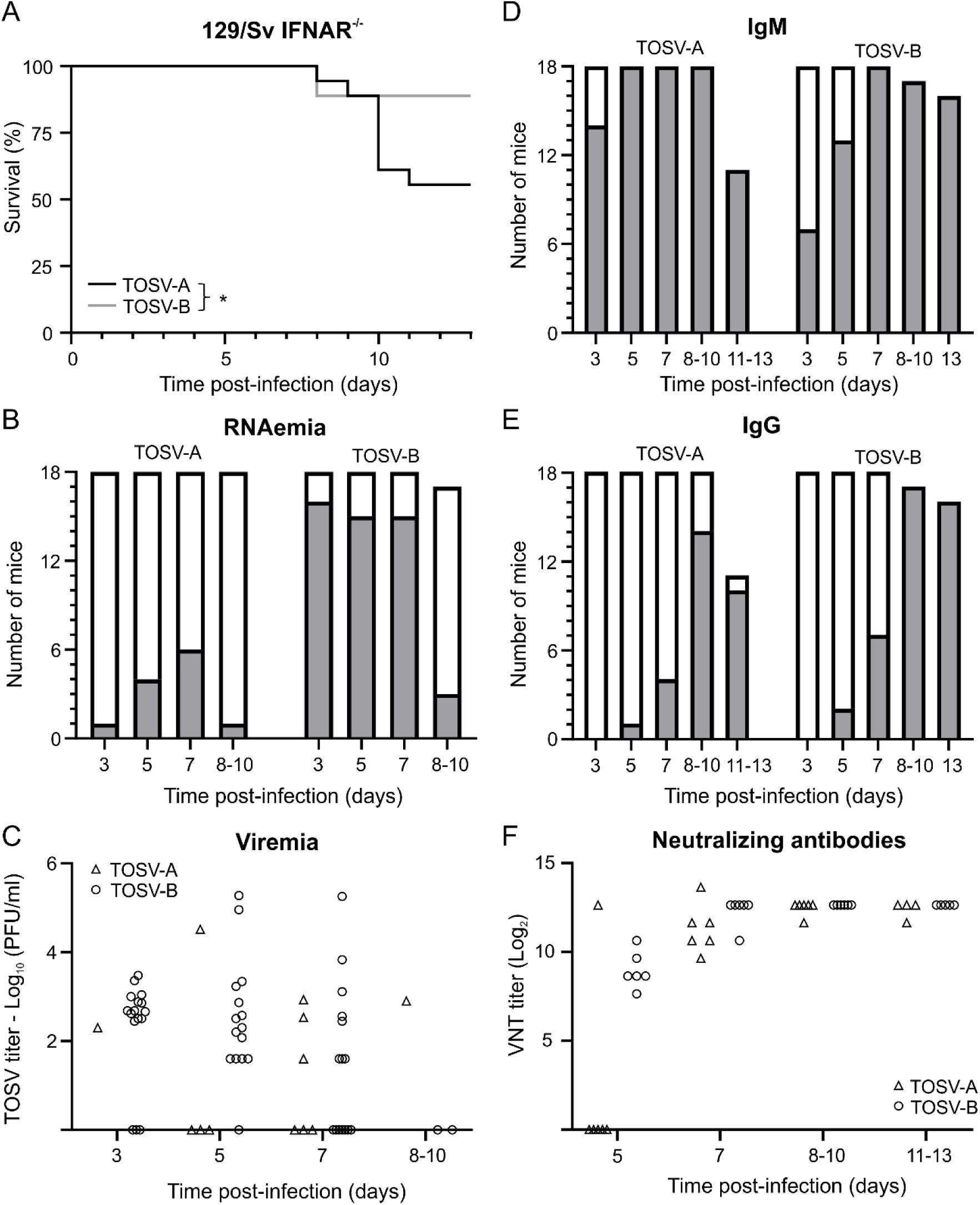
Experimental infection of 129/Sv *ifnar*^-/-^ mice with TOSV-A and TOSV-B. (A) Kaplan-Meier survival curves. 6-8 weeks old female 129/Sv *ifnar*^-/-^ mice (n=18) were infected subcutaneously with 10^3^ PFU of either TOSV-A or TOSV-B. Survival curves were analysed using Log-rank (Mantel-Cox) test. (B) Detection of TOSV RNA in the sera (RNAemia). Sera were collected at 3, 5, 7, and 10 dpi. Levels of viral RNA in sera of infected mice were measured by RT-qPCR targeting segment S. Positive mice sera are represented in gray and negative mice sera are represented in white. Note that only 17 mice were analysed at 8-10 dpi for TOSV-B group because one mouse died suddenly at 8 dpi. Euthanised mice at 8 dpi (one in both TOSV-A and TOSV-B groups) and 9 dpi (one in TOSV-B group) were added to 10 dpi condition (8-10 dpi). (C) TOSV titer in sera (viremia). Viral infectious viral titers were determined by plaque-forming assay for RT-qPCR positive sera. Kinetics of IgM (D) and IgG (E) antibody response against TOSV-A and TOSV-B. Sera were collected at 3, 5, 7, 10 and 13 dpi. Anti-TOSV antibodies within mice sera were detected using in-house IgM and IgG ELISAs. At the indicated day, grey bars represent seroconverted mice and white bars mice with non-detectable anti-TOSV IgM or IgG antibodies Euthanised mice at 8 dpi (one in both TOSV-A and TOSV-B groups) and 9 dpi (one in TOSV-B group) were added to 10 dpi condition (8-10 dpi), when one mouse in TOSV-A group euthanised at 11 dpi was added to 13 dpi condition (11-13 dpi). (F) Virus neutralisation test. Sera (from one independent experiment only, n=6) containing anti-TOSV IgG and/or IgM were tested for the presence of nAbs against TOSV. The neutralizing titre, expressed as log_2_, was determined as the highest dilution of serum that caused complete cytopathic effect.

These data were corroborated by virus titers obtained by plaque-forming assay (Fig 3C). In the sera of TOSV-B-infected mice, mean log_10_ (PFU/ml) values at different time points after infection were similar, although the data distribution was wide, particularly at 5 and 7 dpi. (ranging from 1.6 to 5.3 log_10_ (PFU/ml)). There was a low number of TOSV-A-infected mice showing viremia, however, when TOSV-A-infected mice were viremic, the amount of infectious viral particles detected in the sera was within the same order of magnitude as that observed in TOSV-B-infected mice (Fig 3C). In addition, unlike what we observed in C57BL/6JRj mice, the dynamics of IgM and IgG seroconversion in 129/Sv *ifnar* ^-/-^ mice infected with either TOSV-A or TOSV-B was relatively similar (Fig 2C and 2D, respectively). However, there was a slight delay in the appearance of nAbs in TOSV-A-infected mice. In fact, only one TOSV-A-infected mouse (out of six) produced nAbs at 5 dpi, whereas all TOSV-B-infected mice did (Fig 3F). No other obvious difference was observed at later time points after infection. Overall, our data showed that virulence of TOSV-A in 129/Sv *ifnar* ^-/-^ mice is not directly correlated with viremia, as fewer mice infected with TOSV-A became viremic compared to TOSV-B, but it may be related to a slower production of TOSV nAbs.

### TOSV-A and TOSV-B organ dissemination in 129/Sv *ifnar*^-/-^ mice

To study the dissemination of TOSV-A and TOSV-B, we analyzed by RT-qPCR the presence of TOSV RNA in the brain of mice euthanized during or at the end of the experiment. We showed that all the TOSV-A-infected mice that were moribund and euthanized between 8 and 11 dpi, as well as two mice at 13 dpi, had TOSV RNA in the brain whereas only one TOSV-B-infected mouse was positive at 13 dpi (Fig 4A). Moreover, we detected infectious viral particles with viral titers ranging from 3.5 to 8.4 log_10_ (PFU/g) in mice of TOSV-A group, suggesting that TOSV-A was able to reach and replicate in the brain of 129/Sv *ifnar* ^-/-^ mice, unlike TOSV-B (Fig 4B). Notably, the liver of one TOSV-A-infected mice euthanized at 8 dpi was also positive for TOSV RNA (Ct value = 21.04) and infectious viral particles (1.26 x 10^4^ PFU/g). Interestingly, one of the two TOSV-B-infected mice euthanized at 8 dpi showed an enlarged spleen and we detected both TOSV RNA (Ct value = 20.1) and infectious viral particles (7.7 x 10^2^ PFU/g). Next, as for C57BL/6JRj mice, we investigated TOSV-A and TOSV-B targeted organs in 129/Sv *ifnar* ^-/-^ mice at 3 and 5 dpi. First, we showed that, at 3 dpi, TOSV RNA was detected only in the lungs (two mice out of two) and spleen (1/2) of TOSV-A-infected mice. By 5 dpi, TOSV RNA was also detected in liver (2/2), lungs (2/2), spleen (2/2), and lymph node (1/2) (Fig S1D). Comparatively, most organs of TOSV-B-infected mice were TOSV-RNA positive at both days post-infection. Notably, brain of TOSV-A-or TOSV-B-infected mice were all negatives at 3 and 5 dpi (Fig S1D). Furthermore, we detected viral infectious particles only in spleen (2/2) and lymph nodes (1/1) of TOSV-A-infected mice at 5 dpi, when RT-qPCR data were better correlated with viral titers measured in organs of TOSV-B-infected mice (Fig 4C). Overall, spleen seems to allow active replication of both TOSV strains early after infection, although viral load in this organ at 5 dpi was higher in TOSV-B-infected mice (6.1 x 10^5^ and 2.7 x 10^6^ PFU/g) than in TOSV-A-infected mice (1.7 x 10^3^ and 5 x 10^3^ PFU/g). Altogether, our data showed that TOSV-A can invade and replicate in the brains of half of SC-infected mice whereas TOSV-B spread to most other organs but not the brain.

**Figure 4:**
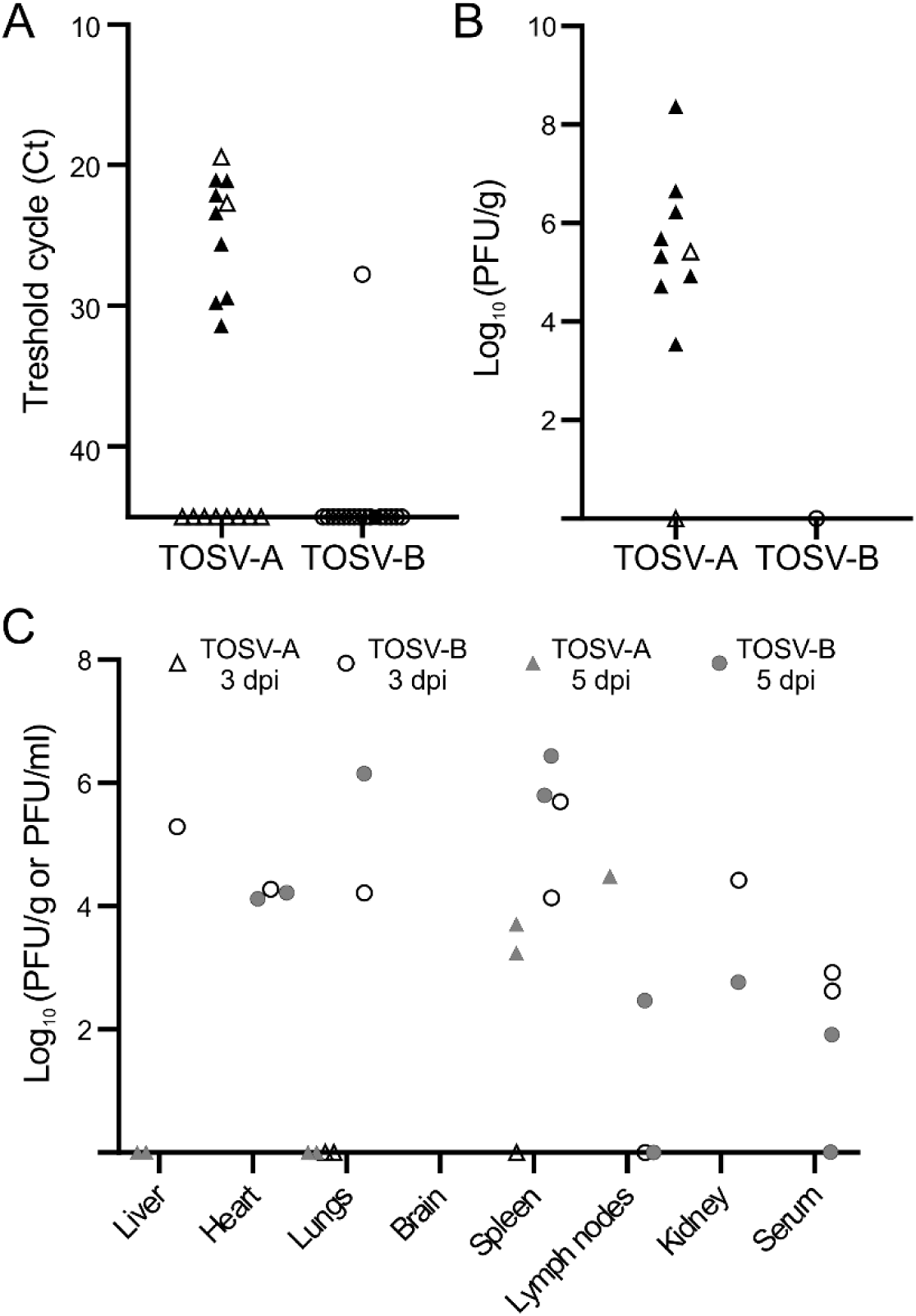
TOSV organ tropism in 129/Sv ifnar^-/-^ mice and viral load in organs/sera. (A) Detection of TOSV RNA in brains of mice euthanised during the experiment (black triangles and circle) or at the end of the experiment (white triangles and circles). Levels of viral RNA were measured by RT-qPCR targeting segment S. (B) Viral loads of TOSV in mouse brains. Viral infectious titers were determined by plaque-forming assay for RT-qPCR positive brains of mice euthanised during the experiment (black triangles) or at the end of the experiment (white triangles and circle). (C) Viral loads of TOSV in mouse organs and serum at 3 and 5 dpi. Mice, subcutaneously infected with 10^3^ PFU of either TOSV-A or TOSV-B, were euthanized at either 3 (white triangles and circles) or 5 dpi (grey triangles and circles). Organs and sera were collected and the presence of TOSV RNA was assessed using RT-qPCR and, if positive, the infectious viral titre (expressed as PFU/g or PFU/ml, respectively) was determined using plaque-forming assay.

### TOSV-B can infect and replicate in mice organotypic brain cultures

Since we did not find infectious viral particles of TOSV-B in the brain, one may hypothesize that this strain is not able to replicate in this organ even if TOSV-B displays higher viral titers in hIPSc-derived neural cells than TOSV-A. To test this hypothesis, we infected OBC isolated from C57BL/6JRj or C57BL/6JRj *ifnar* ^-/-^ mice with 10^3^ PFU of each TOSV strain. In C57BL/6JRj, TOSV-A and TOSV-B reached relatively low but comparable viral titers at 24 (3.6 and 4,4 log_10_(TCID_50_/ml), respectively) and 72 hpi (3.1 and 3.5 log_10_(TCID_50_/ml), respectively) (Fig 5), even though we significantly detected more viral RNA for TOSV-B at both time points (Fig S2). Furthermore, viral titers measured in OBCs isolated from C57BL/6JRj *ifnar* ^-/-^ mice were significantly higher than in OBCs from C57BL/6JRj mice, demonstrating that the IFN response inhibits TOSV replication in OBCs. Interestingly, in this experimental condition, TOSV-B grew to higher titers than TOSV-A at 24 (6.5 and 4,4 log_10_(TCID_50_/ml), respectively) and 72 hpi (9.7 and 7.8 log_10_(TCID_50_/ml) (Fig 5). These data showed that TOSV-B is capable of replicating in the brain of C57BL/6JRj and *ifnar* ^-/-^ mice.

**Figure 5:**
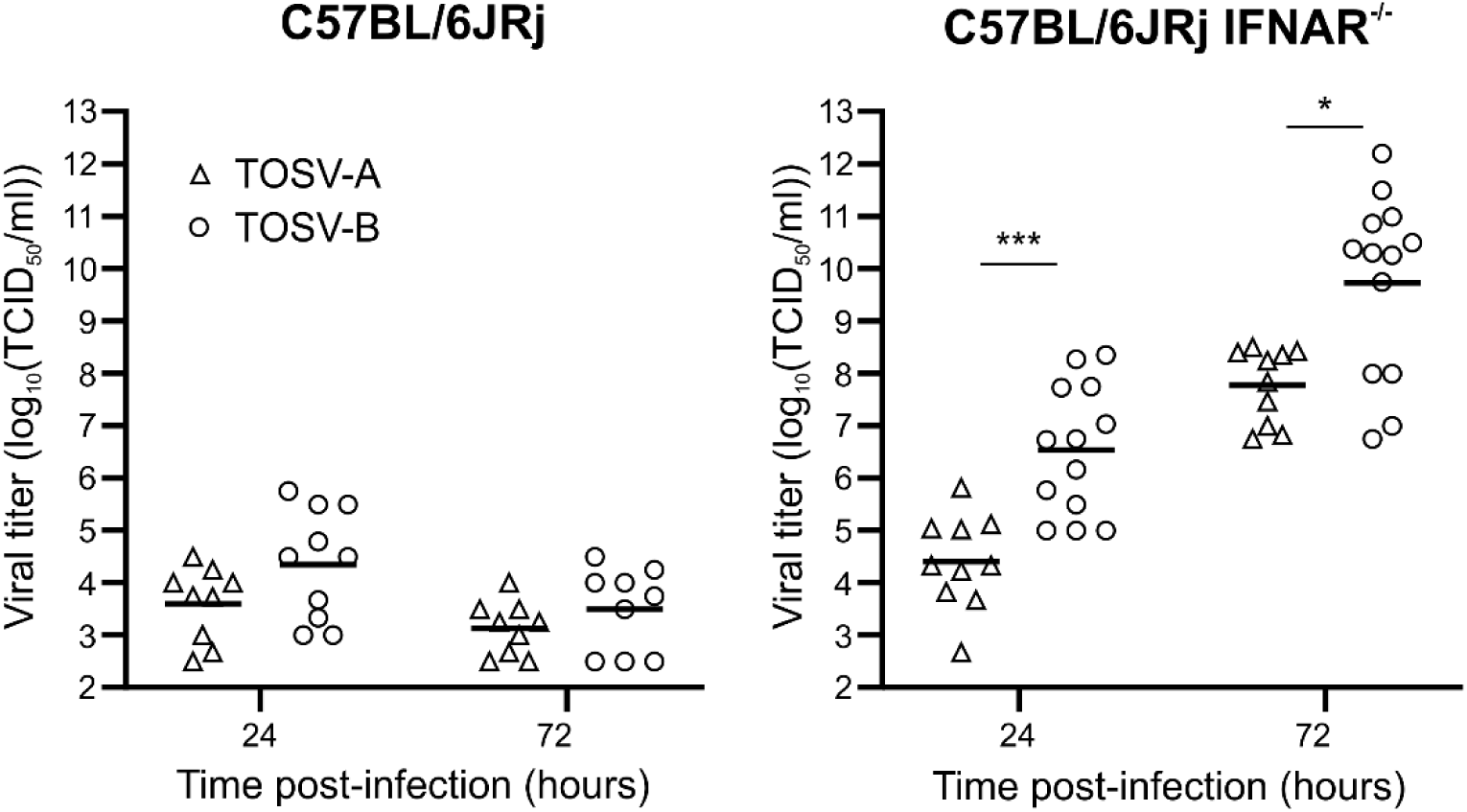
Growth properties of TOSV-A and TOSV-B in mice OBC. Each OBC, obtained from either C57BL/6JRj or C57BL/6JRj *ifnar* ^-/-^ mice, was infected with 10^3^ PFU of either TOSV-A (n=9 and n=10, respectively) or TOSV-B (n=10, 24 hpi -n=9, 72 hpi, and n=13, respectively). Amount of infectious viral particles were determined by endpoint dilution at 24 and 72 hpi. Mean values are represented and comparisons of viral titers at each time point between TOSV-A and TOSV-B were carried out using the Mann-Whitney test; P > 0.05 (ns), P < 0.05 (*), and P < 0.001 (***).

## Discussion

In this study, we characterized two mouse models of TOSV infection. We showed that C57BL/6JRj mice are poorly susceptible to TOSV infection and, mostly, do not develop disease. Importantly, none of the TOSV strains tested were found in the brain of these mice. On the contrary, we observed an increased mortality rate of 129/Sv *ifnar* ^-/-^ mice infected with TOSV, especially with TOSV-A that was detected in the brain of half of infected animals. Interestingly, a previous study reported that *ifnar*^−/−^ mice were susceptible to TOSV strain 1812 (lineage A) infection, with a mortality rate of over 50% of the animals after two weeks post-infection [41]. In addition, brain sections from SC-infected BALB/c mice with 6x10^5^ PFU of the neuro-adapted strain 1812V showed TOSV positivity in endothelial cells of venulae but without apparent disease [42]. These results suggest that IFN response is key to control TOSV infection when inoculated subcutaneously. Nevertheless, *ifnar* ^-/-^ mouse model has already been extensively used to study pathogenesis and virulence of many arboviruses, and our data showed that 129/Sv *ifnar* ^-/-^ mice could be very useful to study the first steps of TOSV infection *in vivo*, dissemination of the virus and TOSV-induced pathogenesis [43].

Both TOSV strains were found in the spleen (5 dpi for TOSV-A and as early as 3 dpi for TOSV-B), and to a lesser extend in lymph nodes where productive infection occurs, confirming that these organs are targeted by TOSV and may constitute a key site of TOSV replication [32]. The earlier and higher viral loads observed in the spleens of TOSV-B-infected mice, compared to TOSV-A, could therefore explain why TOSV-B leads to a higher viremia and spreads more efficiently to several organs (such as liver, heart, lungs, and kidney) compared to TOSV-A. To date, it is still unknown how TOSV-A is spreading to CNS whereas TOSV-B is largely unable to cross the blood-brain barrier, as only one mouse had TOSV-B RNA in its brain, and no detectable viral infectious particles. This cannot be simply explained by an inability of TOSV-B to infect brain cells, as we showed that it replicates even more efficiently than TOSV-A *in vitro* in both hIPSc-derived neural cells and OBCs isolated from C57BL/6JRj *wt* and *ifnar* ^-/-^ mice. Interestingly, given that we showed that all the six TOSV-B-infected mice tested produced nAbs at 5 dpi, when only one TOSV-A-infected mouse did, it is tempting to hypothesize that nAbs response is limiting TOSV-B dissemination to the brain. Especially since it has been suggested that TOSV is freely circulating in the bloodstream (i.e. not cell-associated) [42]. However, we detected TOSV in the brain of the TOSV-A-infected mouse at 13 dpi despite a strong production of nAbs at 5 dpi, indicating that further studies will be required to reveal the pathway by which TOSV-A enters the CNS. TOSV could also infect monocyte-derived dendritic cells (moDCs) and endothelial cells both *in vitro* and *in vivo* [32,42,44,45]. Although we infected 129/Sv *ifnar* ^-/-^ mice that are unable to respond to type 1 IFN, these cells are still able to produce pro-inflammatory cytokines including type 2 and 3 IFN and therefore modulate innate and adaptative immune responses that could differentially impact TOSV replication and dissemination [46,47]. Therefore, comparative studies will be necessary to further explore the replication capacities of both TOSV strains in these cells, as well as their ability to cross the blood–brain barrier using available models and to alter cytokines secretion or antigen-presentation [48]. This is particularly promising since we showed that growth properties of TOSV-A and TOSV-B are markedly different depending on the cell lines tested in this study.

To date, there is no epidemiological evidence that TOSV genetic diversity is impacting its pathogenicity and, to our knowledge, no comprehensive study has compared the biological properties of several strains of TOSV [9]. Here, we showed that two strains belonging to either lineage A or B have their own phenotypic traits. However, it is not yet possible to extend these data to all strains of each genetic lineage, and any attempt to do so would be premature. Moreover, no information is available on the presence of quasispecies in TOSV strains, although this has been shown for other *Phenuiviridae* such as Rift Valley fever virus with consequences on its virulence [49]. Here, both TOSV-A and TOSV-B strains were not plaque purified which means that the virus stocks used in this study may contain viral quasispecies or defective interfering viruses with potential effect on their viral replication and/or dissemination. The recent development of a reverse genetic systems for TOSV will certainly be key to broaden the scope of the results obtained in this study by identifying, for example, the viral determinants responsible for the efficient dissemination of TOSV-A to CNS or the high viremia of TOSV-B in 129/Sv *ifnar* ^-/-^ mice [18,20].

Taken together, our results demonstrates that 129/Sv *ifnar* ^-/-^ is a relevant mouse model that offers new perspectives to shed some light of TOSV-induced pathogenesis. Our study also highlight the importance to further study the role of genetic diversity of TOSV in its virulence, organ tropism and vector transmission in order to better understand the epidemiology of this neglected sand fly-borne pathogen.

## Acknowledgments

The authors would like to thank Justine Girard, Etienne Ossona de Mendez, Marie-Pierre Confort and Barbara Viginier for their implication in this work and useful discussions. TOSV strain MRS2010-4319501 was obtained from the European Virus Archive goes Global (EVAg) project that has received funding from the European Union’s Horizon 2020 research and innovation program under grant agreement No 653316.

## Authors contribution

Conceptualization and investigation: Philippe Marianneau, Frédérick Arnaud and Maxime Ratinier

Data curation and formal analysis: Marlène Roy, Sandra Lacôte, Cyrille Mathieu, Bertrand Pain, Philippe Marianneau, Frederick Arnaud and Maxime Ratinier

Methodology: Marlène Roy, Sandra Lacôte, Sophie Desloire, Adrien Thiesson, Coralie Pulido, Noémie Aurine, Cyrille Mathieu, Bertrand Pain and Maxime Ratinier

Resources: Cyrille Mathieu, Bertrand Pain, Philippe Marianneau and Maxime Ratinier

Supervision and validation: Cyrille Mathieu, Bertrand Pain, Philippe Marianneau, Frédérick Arnaud and Maxime Ratinier

Writing – original draft: Maxime Ratinier

Writing – review & editing: Marlène Roy, Sandra Lacôte, Sophie Desloire, Adrien Thiesson, Coralie Pulido, Noémie Aurine, Cyrille Mathieu, Bertrand Pain, Philippe Marianneau, Frédérick Arnaud, Maxime Ratinier

