## Supplemental_Figures_S1_S2 for "Two strains of Toscana virus show different virulence and replication capacity in mice and cell culture models"

### Supporting information

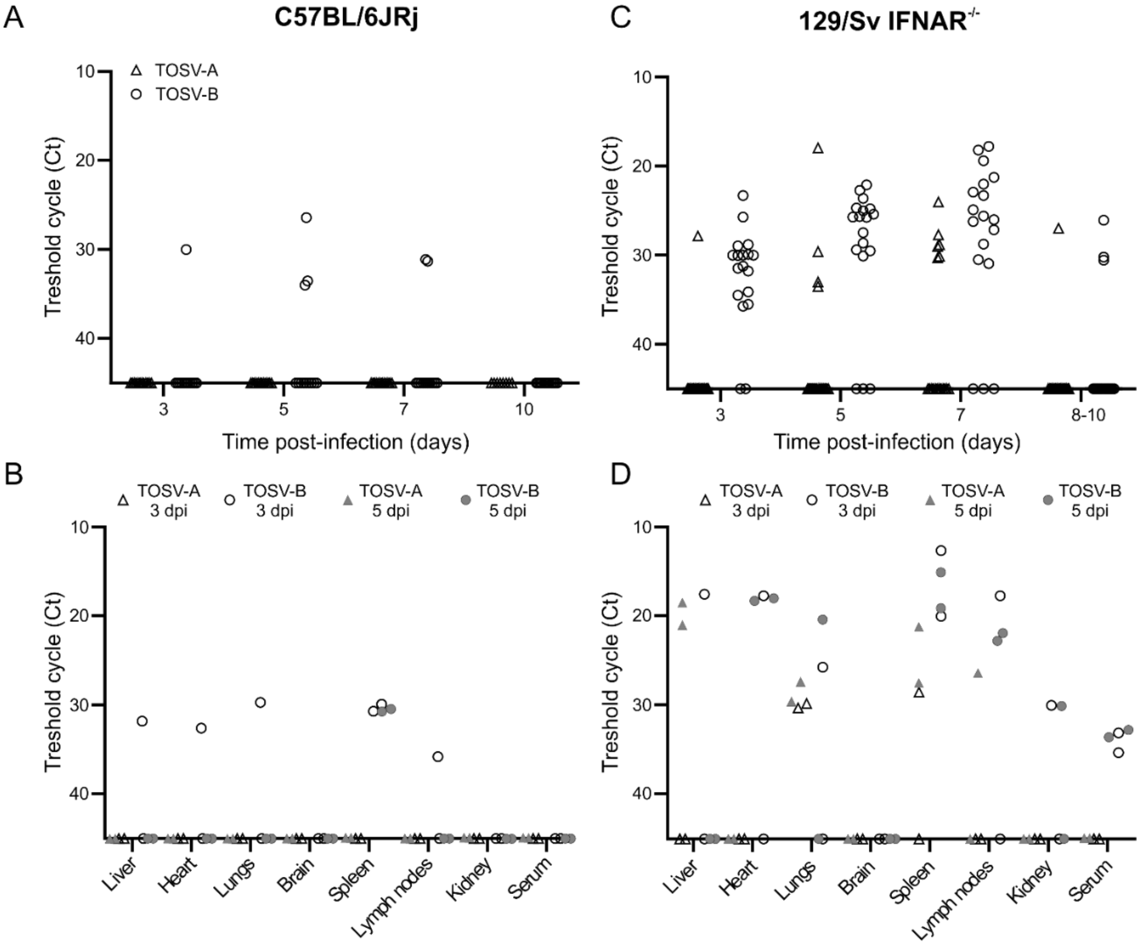


**<u>Figure S1:</u> Levels of TOSV RNA in sera and organs of C57BL/6JRj and 129/Sv *ifnar$^{-/-}$* mice.** (A) Detection of TOSV RNA in the sera (RNAemia) of C57BL/6JRj mice. Sera were collected at 3, 5, 7, and 10 dpi. Levels of viral RNA in sera of infected mice were measured by RT-qPCR targeting segment S. Cycle threshold (Ct) values are represented. (B) Levels of viral RNA in the organs and serum of C57BL/6JRj mice at 3 and 5 dpi. Mice, subcutaneously infected with $10^3$ PFU of either TOSV-A or TOSV-B, were euthanized at either 3 (white triangles and circles) or 5 dpi (grey triangles and circles). Organs and sera were collected and the presence of TOSV RNA was assessed using RT-qPCR targeting segment S. Ct values are represented. (C) Detection of TOSV RNA in the sera (RNAemia) of 129/Sv *ifnar$^{-/-}$* mice. Sera were collected at 3, 5, 7, and 10 dpi. Levels of viral RNA in sera of infected mice were measured by RT-qPCR targeting segment S. Ct values are represented. (D) Levels of viral RNA in the organs and serum of 129/Sv *ifnar$^{-/-}$* mice at 3 and 5 dpi. Mice, subcutaneously infected with $10^3$ PFU of either TOSV-A or TOSV-B, were euthanized at either 3 (white triangles and circles) or 5 dpi (grey triangles and circles). Organs and sera were collected and the presence of TOSV RNA was assessed using RT-qPCR targeting segment S. Ct values are represented.

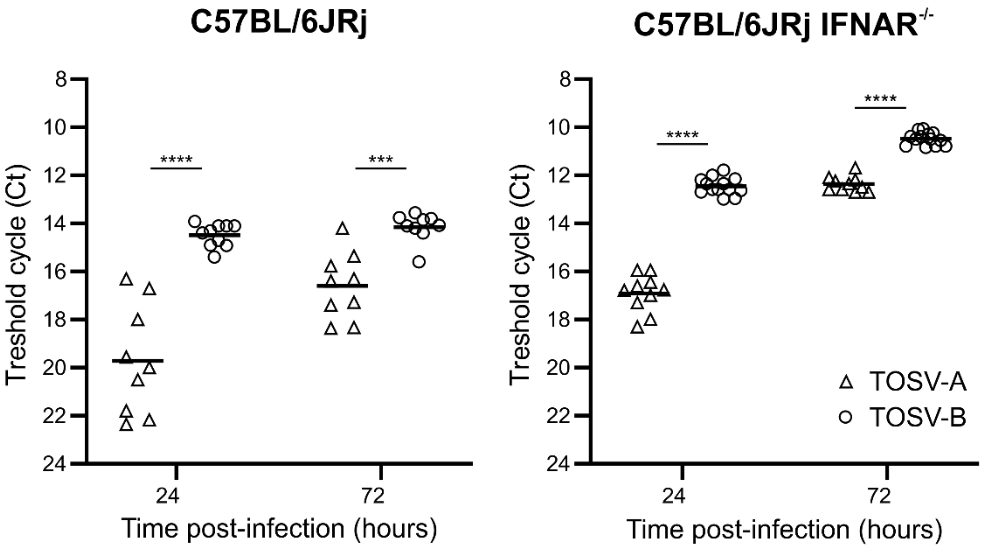


**Figure S2:** **Levels of TOSV RNA in OBC.** Each OBC, obtained from either C57BL/6JRj or C57BL/6JRj *ifnar* $^{-/-}$ mice, was infected with $10^3$ PFU of either TOSV-A or TOSV-B. Levels of viral RNA in each OBC were measured by RT-qPCR targeting segment S. Individual Ct and mean values are represented. Comparisons of Ct values at each time point between TOSV-A and TOSV-B were carried out using the Mann-Whitney test; $P < 0.001$ (***) and $P < 0.0001$ (****).
